# Temperature and frequency dependence of conduction along sympathetic preganglionic axons

**DOI:** 10.64898/2026.05.20.726598

**Authors:** Mallika Halder, Shawn Hochman

## Abstract

Sympathetic preganglionic neurons (SPNs) distribute signals widely across paravertebral ganglia, yet the reliability of spike propagation along their predominantly unmyelinated axons remains poorly defined. We examined temperature- and activity-dependent modulation of SPN axonal conduction using an ex vivo adult mouse thoracic sympathetic chain preparation. Population compound action potentials (CAPs) were evoked by supramaximal stimulation of T10 ventral roots and recorded from branching axons in interganglionic compared to unbranching axons in the splanchnic nerve.

At physiological temperature (36°C), scaled CAP magnitude was reduced by ∼50% relative to 22°C, with preferential loss of slower-conducting axonal components. These reductions are consistent with substantial temperature-dependent decreases in effective axonal recruitment, likely reflecting conduction failure in a large fraction of SPNs. Losses were more pronounced in interganglionic pathways, suggesting increased vulnerability in branching projections.

To assess activity-dependent effects, stimuli were delivered at 1, 5, and 20 Hz with focus on 5 and 20 Hz stimulus trains (20s duration). The overall time-course of train-evoked depression was similar across temperatures; however, the underlying axonal populations differed. At 22°C, slower-conducting axons exhibited marked frequency-dependent depression, whereas at 36°C the remaining faster-conducting axons displayed facilitation, particularly at 20 Hz. Slower-conducting responses also showed post-train potentiation at physiological temperature.

These findings indicate that SPN axonal conduction is not uniformly reliable and is strongly modulated by temperature and activation history. Preferential vulnerability of slow-conducting, likely small-diameter and branching axons identifies axonal conduction as a physiologically regulated site of gain control in sympathetic output.

## INTRODUCTION

Sympathetic preganglionic neurons (SPNs) distribute signals widely across paravertebral ganglia, providing the primary drive to postganglionic sympathetic neurons that regulate cardiovascular, thermoregulatory, and metabolic functions. These projections are typically assumed to transmit action potentials reliably, such that preganglionic firing is faithfully translated into postganglionic output (McLachlan, 2003). However, this assumption has not been rigorously tested under physiological conditions. Given the distinctive structural features of SPN axons, including their predominantly unmyelinated nature and extensive branching within the sympathetic chain, it is unclear whether spike propagation remains reliable across their full anatomical extent.

Axonal geometry and diameter are key determinants of conduction reliability. SPN axons in mouse are overwhelmingly unmyelinated and extremely small in caliber, with mean diameters on the order of 0.4 μm and branch diameters as small as 0.1 μm (Lewis and Burton, 1977). Such small-diameter axons operate with a reduced safety factor for propagation and are therefore particularly sensitive to perturbations in membrane conductance and ionic currents. In other systems, axonal branch points are well established as sites of conduction failure, especially when the available depolarizing current is insufficient to support propagation into multiple daughter branches (Levy, 1980; Debanne et al., 2011). These considerations suggest that SPN axons, by virtue of their size and branching architecture, may be especially vulnerable to failures of spike propagation, with potential consequences for the effective distribution of sympathetic signals.

Temperature is a critical determinant of axonal excitability and conduction reliability, particularly in unmyelinated fibers. Increases in temperature accelerate channel kinetics but can reduce spike amplitude and duration, thereby lowering the safety factor for propagation (Hodgkin and Katz, 1949; Paintal, 1965; Pekala et al., 2016). Conversely, cooling can broaden action potentials and may enhance conduction reliability. Despite this, prior physiological studies of multisegmental SPN signaling have been performed at room temperature (Blackman and Purves, 1969; Nja and Purves, 1977; Lichtman et al., 1979, 1980; Purves and Lichtman, 1980), raising the possibility that the reliability and extent of SPN signal distribution under physiological conditions have been substantially mischaracterized. In addition to temperature, activation history is known to influence conduction in unmyelinated axons through activity-dependent hyperpolarization and sodium channel inactivation, which can lead to conduction slowing and failure during repetitive firing (Renganathan et al., 2002; De Col et al., 2008; Obreja et al., 2010; Rama et al., 2018).

Here, we tested the hypothesis that spike propagation in SPN axons is not uniformly reliable and is strongly modulated by temperature and activation history. Using an ex vivo adult mouse preparation that preserves the thoracic sympathetic chain and ventral roots, we recorded population compound action potentials (CAPs) evoked by supramaximal stimulation of SPN axons. By comparing responses at room and physiological temperatures and across different stimulation frequencies, we assessed how temperature and activation history influence effective axonal recruitment. We further examined whether slower-conducting axons, which likely correspond to smaller diameter and more extensively branching fibers, are preferentially vulnerable to conduction failure. These experiments were designed to determine whether axonal conduction reliability represents a previously unrecognized site of gain control in sympathetic output.

## METHODS

### Multisegmental *ex vivo* paravertebral preparation

Adult (8 weeks+) C57Bl/6 mice of both sexes were anesthetized with ketamine (100mg/kg) and xylazine (10mg/kg) mix by intraperitoneal injection, following light anesthetization in an isoflurane chamber. The thoracic vertebral column and adjacent ribs are excised and transferred to a Sylgaard dish containing ice cold, oxygenated (95% O_2_ / 5% CO_2_) high-Mg^2+^/low-Ca^2+^ artificial cerebral spinal fluid (aCSF) containing (in mM), [NaCl 128, KCl 1.9, MgSO_4_ 13.3, CaCl_2_ 1.1, KH_2_PO_4_ 1.2, glucose 10, NaHCO_3_ 26]. Following a complete laminectomy and vertebrectomy, the spinal cord (SC) and dorsal roots (DR) are exposed and removed. The remaining thoracic chain ganglia, in continuity with communicating rami, spinal nerves and ventral roots are cleaned of excess fat and muscle. The tissue is transferred to a Sylgaard recording chamber superfused with oxygenated aCSF containing (in mM), [NaCl 128, KCl 1.9, MgSO_4_1.3, CaCl_2_2.4, KH_2_PO_4_ 1.2, glucose 10, NaHCO_3_ 26], at ∼40ml/minute at 28C and allowed to rest for 1 hour.

### Electrophysiology

#### Temperature and activity-dependence experiments

Glass recording electrodes were placed on interganglionic nerve at the caudal cut end of the T12 ganglion and the adjacent splanchnic nerve using ‘trumpet’ shaped electrodes with progressively reduced internal diameter reaching ∼100-125 μm (Halder et al., 2021). Glass suction recording electrodes were positioned on the T10 ventral root for stimulation (200-250 *µ*m tip diameter). The distance between stimulation a recording sites was 5 mm (**Fig. 1A**). For temperature experiments, bath temperature was maintained at 36°C by a Peltier device (built in-house by Mr. Bill Goolsby) prior to start of experiment. The experimental protocol involved delivery of 5 separate stimulus trains at 60 second intervals. At this interval evoked responses recovered to pre-train values. The design involved data capture of baseline for two seconds prior to delivery of a pre-train stimulus occurring 1 before the 20 second stimulus train (5 Hz or 20 Hz), then followed by a post-train stimulus delivered two seconds later to assess history-dependent changes (see **Fig. 3A**). Quantified are (A) responses at train start, (B) 1 second into the train, (C) the last evoked response in the train, and (D) 2 seconds post-train stimulation. During the experiment, the bath was then cooled to 22°C to assess temperature-dependent changes in preganglionic conductance. The tissue was allowed to stabilize for at least 15 minutes before data acquisition. All recorded data were digitized at 50 kHz (Digidata 1322A 16 Bit DAQ, Molecular Devices, U.S.A.) with pClamp acquisition software (v. 10.7 Molecular Devices). Recorded signals were amplified (5000x) and low pass filtered at 3 kHz using in-house amplifiers.

**Figure 1.**
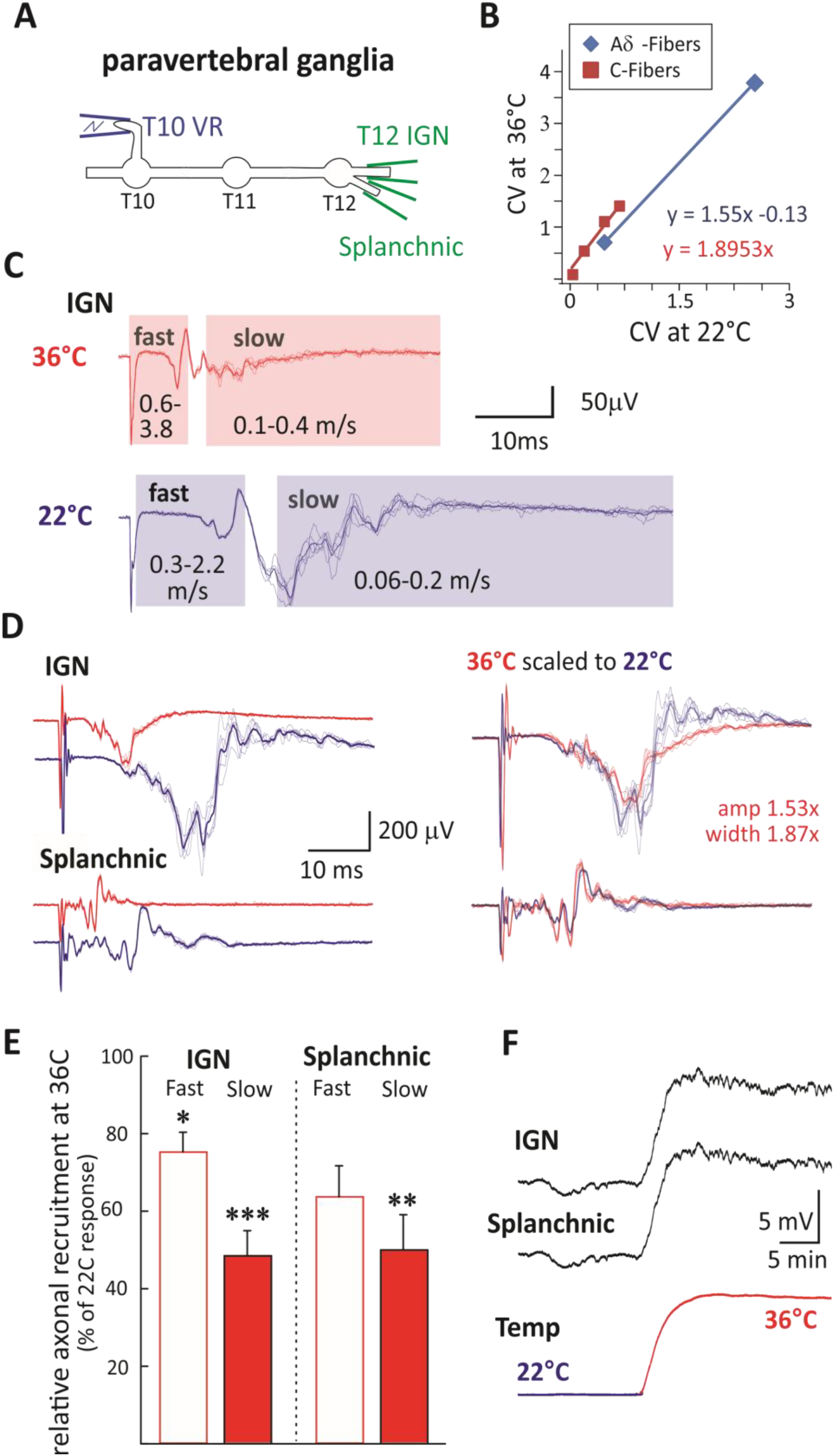
Increased temperature leads to reduced axonal recruitment. **[A]** Experimental setup showing SPN electrical stimulation in the T10 ventral root (VR) while recording evoked CAPs from the caudal T12 IGN and the splanchnic nerve. **[B]** Graph derived from Table 1 in Pinto et al (2008), estimating the changes in conduction velocity for Aδ and C-fibers, which exhibit conduction velocities comparable to SPN fibers observed in mice. Using trend line formulas and experimental data collected at 22°C (fast: 0.3-2.2 m/s; slow: 0.06-0.2 m/s), predicted changes to conduction velocity ranges were extrapolated for fast (0.6-3.8 m/s) and slow populations (0.1-0.4 m/s) at 36°C as shown in temp and frequencies comparison studies summarized in Table 1. **[C]** Example T° differences in CAP appearance and separation into fast and slow groups. Fast and slow SPNs within conduction velocity ranges above are highlighted with boxes. **[D]** Scaling approach shown in another example recording. Left panels show difference in responses at 36°C and 22°C. Right panels show overlaid volleys with 36°C response amplified using pre-determined scale factors (shown). Note preferential loss of slower responses. **[E]** Evoked responses from 36°C and 22°C were rectified, integrated, scaled, and compared. The percent reduction in magnitude of evoked response was determined for fast and slow-conducting epochs of the CAP. The IGN slower-conducting axons exhibited a substantially higher % reduction in evoked response (48.5±6.4%) compared to that seen in faster-conducting axons (75.3±5.1%; p<0.05) [n=6]. For the splanchnic nerve these values were reduced by 50.0±9.1% and 63.7±8.0%, respectively [n=5]. (*, p<0.05; **p<0.01***; p<0.001; paired t-tests). **[F]** Example extracellular DC recordings of changes in polarization in both IGN and splanchnic with temperature increases from 22°C to 36°C consistent with intracellular membrane hyperpolarization ([n=3] 7.5±1.2 and 7.1±1.1 mV, respectively).

### Pharmacology

As recordings of population axonal spiking could include synaptically-mediated responses, we block synaptic activity with the nicotinic acetylcholine receptor antagonist hexamethonium bromide (100 μM) (Mason, 1962) and included pancuronium (20 µM) to prevent possible intercostal muscle activity (both from Sigma-Aldrich).

**Table 1.** Conduction velocity range (in m/s) of thoracic SPN axons at 22° and 36°C.

|  | 22°C | 36°C |
| --- | --- | --- |
| IGN (n=6) | 0.10±0.001 - 0.40±0.04 | 0.19±0.02 - 0.81±0.06 |
| Splanchnic (n=5) | 0.12±0.02 - 3.1±1.5 | 0.21±0.03 - 7.0±3.9 |

### Analysis

Evoked SPN CAPs, were recorded following VR stimulation once every 60 seconds. Supramaximal recruitment of slow-conducting axons was achieved with electrical stimuli delivered at 200µA, 500µs, as longer pulse durations ensure recruitment of unmyelinated axons (Thompson et al., 1990; Brocker and Grill, 2013). Recorded CAPs were divided into CV-subpopulations depending on latency of arrival from time of stimulation. For each CV population, 5 episodes were rectified and integrated using Clampfit software. At this point background noise was accounted for by subtracting an equal duration of baseline noise to provide a singular, comprehensive measure of CAP size. All 5 episodes were then averaged to ensure a more accurate representation of the CAP. In the analysis of T° effects, we employed paired t-tests to evaluate the significance of differences between T° as well as between CAP epochs within each nerve. One-way repeated measures ANOVA was used for tests across multiple groups. For post-hoc detailed pairwise comparisons, a Bonferroni t-test analysis was utilized. Values are reported as mean percent of baseline ± standard error (**SEM**). Stimulus trains at 5 and 20 Hz (20s duration) were undertaken. Comparison of total amplitude changes during trains at 20 Hz assessed a shorter response interval as a component of the slower response was interrupted by the subsequent stimulus artifact in the train. Quantification of evoked responses were commonly separated into slower and faster conducting components whose range differed at 22°C and 36°C (**Table 1**; also see **Fig. 1B**).

## RESULTS

### Temperature-dependent reduction in population recruitment of SPN axons

We first examined the effect of temperature on SPN axonal conduction by comparing evoked compound action potentials (CAPs) recorded at room temperature (22°C) and physiological temperature (36°C). Conduction velocities increased with temperature, approximately doubling between 22°C and 32°C (**Table 1**), consistent with established temperature dependence of axonal conduction.

To compare responses across temperatures, CAPs recorded at 36°C were scaled to account for temperature-dependent changes in conduction velocity, spike amplitude, and duration. Utilizing the framework provided by Pinto et al 2008 (Pinto et al., 2008), which details T°-dependent changes in nerve fiber signal speeds, we adjusted the spike CV from 36°C to 22°C (**Fig. 1B**,**C**). Then, we scaled the amplitude and width by factors of 1.53 and 1.87, respectively, corresponding to the T° decrease from 36°C to 22°C, using reference data (Hodgkin and Katz, 1949; Stegeman and De Weerd, 1982) (**Fig. 1D**). In trying to fit our data using these amplitude and width values, we observed that while width values were accurate, scaling by 1.53 generally over amplified values but used these values nonetheless so that if anything changes observed would be underestimated. Following scaling, CAP magnitude at 36°C was significantly reduced relative to 22°C in branching axons in the interganglionic (IGN) and in unbranching axons in splanchnic nerve recordings. In the IGN, responses were reduced to 48.8 ± 6.4% of scaled 22°C values (p < 0.001, n = 6), and in the splanchnic nerve to 52.8 ± 7.9% (p < 0.01, n = 5). Overall, these data demonstrate a substantial increase in conduction failures at physiological temperature.

### Preferential loss of slower-conducting axonal populations

To determine whether temperature-dependent reductions in CAP magnitude were distributed uniformly across axonal populations, responses were separated into fast- and slow-conducting components based on conduction latency (**Table 2**).

**Table 2.** Conduction time ranges from start of stimulus used to quantify magnitude changes in fast and slow conducting axons at 22° and 36°C.

| T° & category | Conduction velocity range | Epoch duration |
| --- | --- | --- |
| 22°C fast | 2.2-0.31 m/s | 2.0-14 ms from stim. onset |
| 22°C slow | 0.23-0.06 m/s | 19-70 ms from stim. onset |
| 36°C fast | 3.4-0.58 m/s | 1.3-7.6 ms from stim. onset |
| 36°C slow | 0.44-0.11 m/s | 10-40 ms from stim. onset |

At 36°C, reductions in CAP magnitude were significantly greater in slower-conducting components compared to faster-conducting components. In the IGN, slower-conducting responses were reduced to 48.5 ± 6.4% of scaled 22°C values, whereas faster-conducting responses were reduced to 75.3 ± 5.1% (p < 0.05; **Fig. 1E**). Similar trends were observed in the splanchnic nerve (slow: 50.0 ± 9.1%; fast: 63.7 ± 8.0%). These findings indicate that conduction failures are dominated by preferential loss in slower conducting axons at physiological temperatures.

### Maximal recruitment and membrane potential changes with temperature

We next assessed whether changes in axonal recruitment could be attributed to altered stimulus evoked recruitment. In other experiments (Halder et al., 2026) we determined that a stimulus magnitude of 200μA, 500μs was suprathreshold for axonal population recruitment and used this intensity in the current study. Comparing incidence of maximal recruitment in separate studies where higher stimulus intensities were explored at 22°C vs 32-36°C we observed that 200μA, 500μs was suprathreshold or near maximal recruitment in 8/10 experiments at 22°C and 12/13 experiments at 32-36°C. indicating that maximal recruitment levels were similar across temperatures.

In comparison, direct current (DC) recordings revealed temperature-dependent membrane polarization changes. Increasing temperature from 22°C to 36°C produced a DC depolarization of 7.5 ± 1.2 mV in the IGN and 7.1 ± 1.1 mV in the splanchnic nerve (n = 3) consistent with membrane hyperpolarization (**Fig. 1F**). Such changes are consistent with a change in membrane conductance at higher temperatures.

### Frequency-dependent modulation of CAP magnitude at 22°C

We next examined the effects of activation history on SPN conduction using stimulus trains delivered at 5 Hz and 20 Hz.

At 22°C, CAP magnitude exhibited frequency-dependent depression during stimulus trains in both IGN and splanchnic recordings (**Fig. 2**). Depression increased with stimulus frequency and stabilized by the 5th–6th pulse.

**Figure 2.**
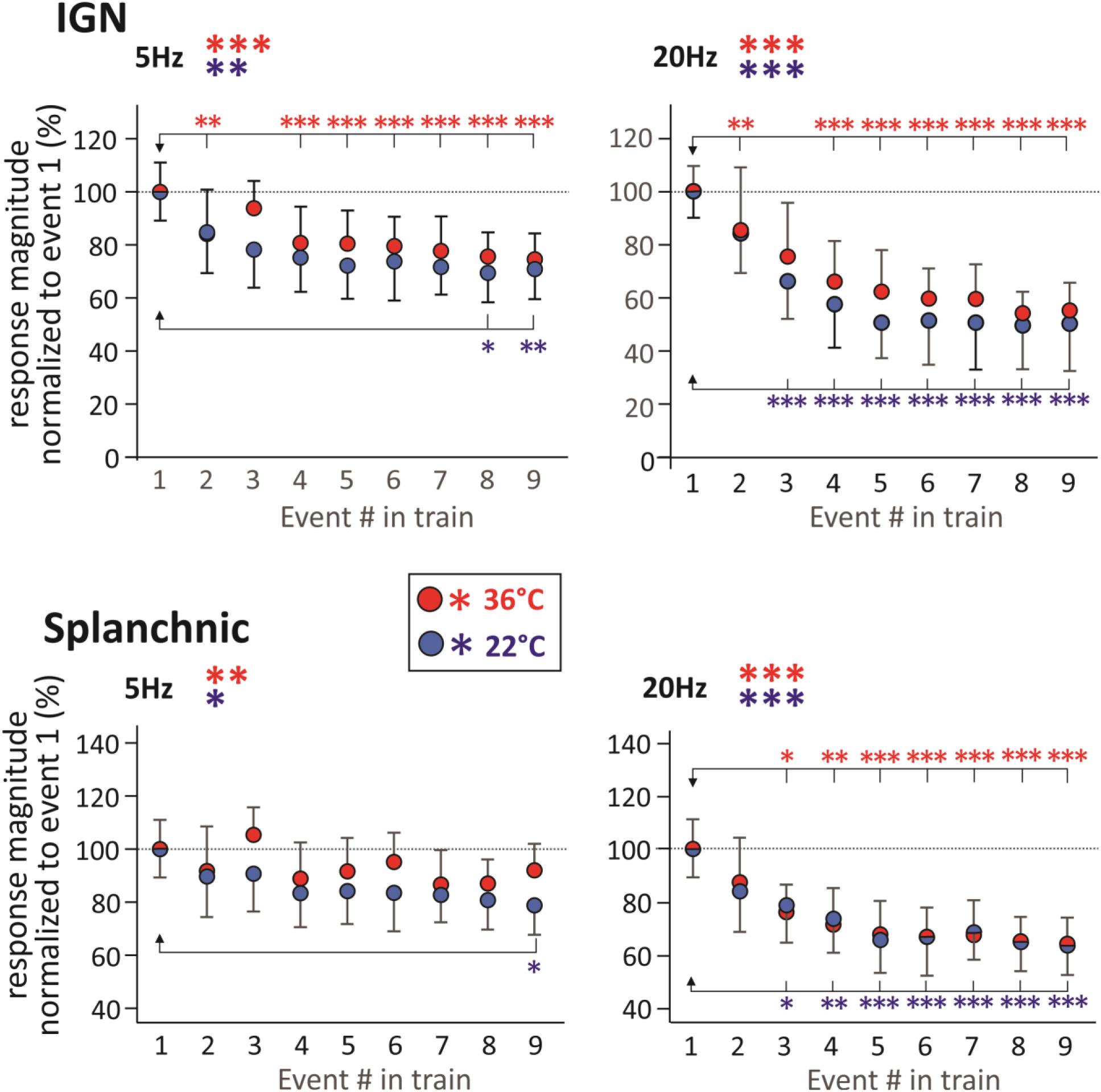
Effect of pulse number on change in evoked CAP response amplitude during 5Hz and 20Hz stimulation at 22° and 36°C. Percentage relative changes in CAP responses relative to the 1^st^ evoked response in the train are compared between 22°C and 36°C at 5 and 20Hz in IGN and splanchnic nerves. There was a significant effect of activation history during the train (events 1-9) in all conditions in both nerves [n=4] (*, p<0.05; **, p<0.01, **; and p<0.001, respectively; RM ANOVA). Relative changes were comparable within nerves at both temperatures and frequencies. Response depression accounted for all significant pairwise response changes in the train relative to the first response (Bonferroni t-test). Note that responses tended to stabilize by the 5-6th pulse.

When then analyzed by axonal subpopulations at defined epochs during the train as depicted in **Fig. 3A**). At 22°C slower-conducting components exhibited pronounced reductions in CAP magnitude during trains, whereas faster-conducting components were largely unaffected (**Fig. 3B**). For example, in the IGN at 5 Hz, slower-conducting responses were reduced to 40.2 ± 4.7% and 29.1 ± 8.0% of baseline at 1s within and at the end of the train (epochs B and C, respectively; p < 0.05 and p < 0.001), while faster-conducting responses did not differ significantly from baseline (**Fig. 3B**_**1**_). Similar patterns were observed at 20 Hz, where slower-conducting IGN responses were reduced to 25.7 ± 10.4% at the end of the train (epoch C; p < 0.001; **Fig. 3B**_**2**_).

**Figure 3.**
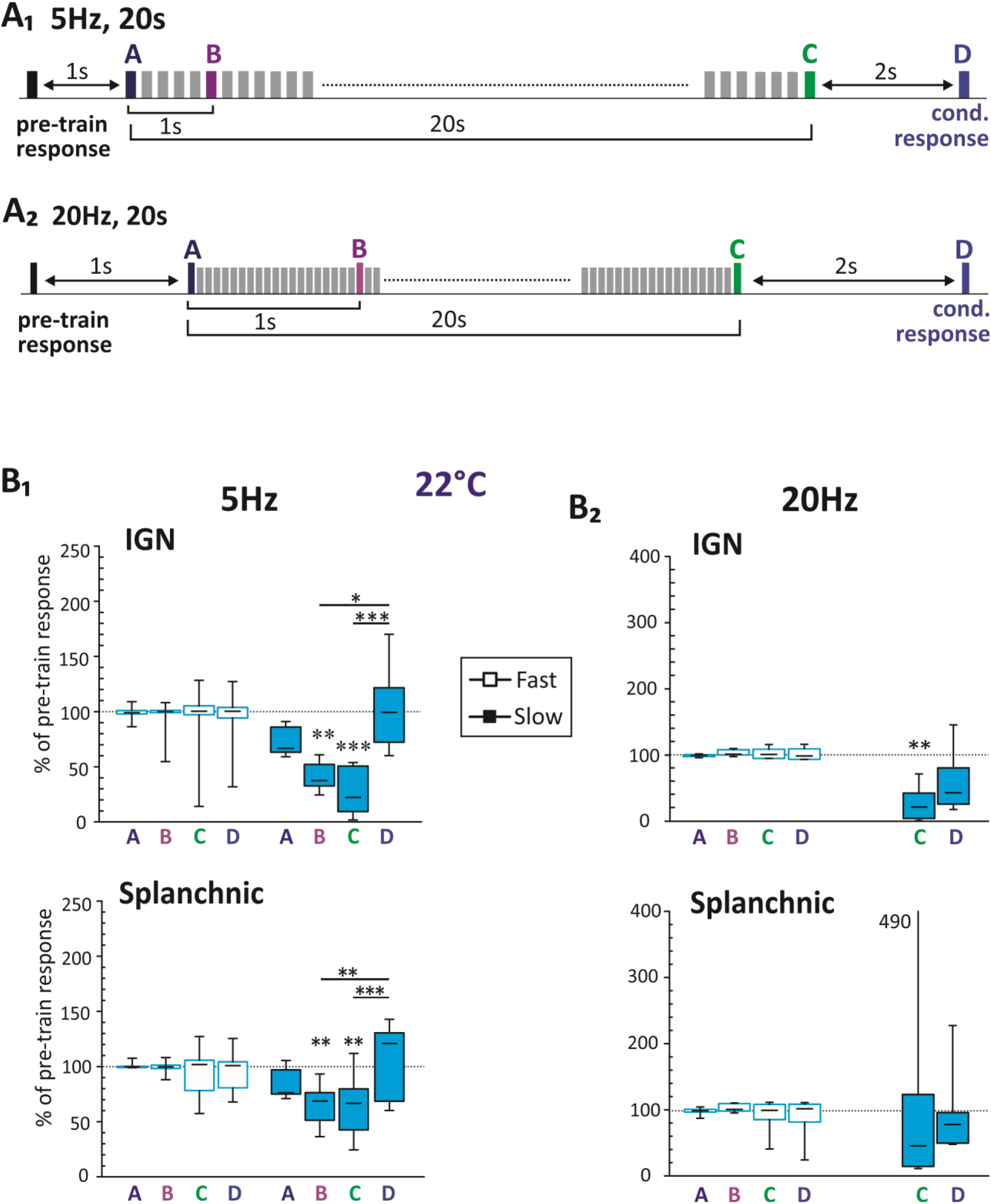
Effect of 5Hz and 20 Hz trains on evoked response at 22°C. **[A_1_]** Experimental design: 5Hz, 20 sec stimulus train initiated 1 sec after the pre-train (control) response. Quantified are responses at train start (A), 1 second into the train (B), the last evoked response in the train (C), and 2 seconds post-train stimulation to assess history-dependent changes (D). The responses evoked for each epoch were divided into faster and slower conducting components. Responses were analyzed against the initial pre-train response for potential history-dependent effects. The rectified integral values of both faster- and slower-conducting CAP components from these epochs are presented as percentages of the rectified-integral values of the baseline pre-train response. We particularly evaluated epochs C & D history-dependent effects observed following prolonged stimulus train delivery (20s). **[A_2_]** Experimental design: 20Hz, 20 sec stimulus train. **[B_1_]** 5 Hz. Faster axon CAPs were unaffected by the 5Hz stimulation train in both IGN and splanchnic nerves. Slower axons exhibited a transient decrease in CAP magnitude during the train, with a subsequent recovery after 2 seconds. Specifically, slow IGN axons displayed a reduction to 40±5% and 29±8% in epochs B and C, respectively, while splanchnic slow axons demonstrated a consistent depression to 66±7% and 66±10% in the same epochs. **[B_2_]** 20 Hz. At 22°C, faster axons in both IGN and splanchnic nerves showed no significant changes, but slower conducting fibers were more variable and underwent significant depression in the IGN nerve (C) at 26±10%. The conditioned pulse (Epoch D) saw a recovery in splanchnic slow axons but remained diminished at 50±17% for IGN (p<0.01). Note: slow-conducting units from Epochs A-B were excluded in the 20Hz stimulation analysis because the CAP arrival was slower than the subsequent pulse in the train. Statistical significance was assessed via repeated measures ANOVA and post hoc tests where relevant ([n=7];*, p<0.05; **, p<0.01, **; and p<0.001, respectively).

As we were unable to selectively quantify the slower-conducting volley components during the 20 Hz train due to overlap with stimulus artifact, comparison of ‘pre-train’ evoked responses are to the final response in the pulse train (epoch C). A summary of results obtained at 22°C is provided in **Table 3**.

**Table 3.** 22C results of stimuli delivered at 5 & 20 Hz for 20 seconds (% of control). (*, p<0.05; **, p<0.01, **; and p<0.001, respectively)

| N=7 | Fast SPNs (0.31-2.2 m/s) |  | Slow SPNs (0.06-0.23 m/s) |  |
| --- | --- | --- | --- | --- |
|  | 5Hz | 20Hz | 5Hz | 20Hz |
| **IGN** |  |  |  |  |
| [A] | 98.9 ± 2.5 | 99.4 ± 0.8 | 71.7 ± 4.6 |  |
| [B] | 94.8 ± 6.8 | 103.3 ± 18.6 | 40.2 ± 4.7* |  |
| [C] | 92.2 ± 13.6 | 102.5 ± 3.3 | 29.1 ± 8.0*** | 25.7 ± 10.4** |
| [D] | 94.0 ± 11.1 | 101.4 ± 3.6 | 102.8 ± 13.6 | 56.2 ± 19.1 |
| **Spl.** |  |  |  |  |
| [A] | 101 ± 1.1 | 97.9 ± 2.0 | 84.5 ± 5.0 |  |
| [B] | 99.2 ± 2.2 | 102.2 ± 2.2 | 65.7 ± 6.6* |  |
| [C] | 96.1 ± 8.4 | 92.4 ± 9.2 | 65.5 ± 10.4* | 109.7 ± 65.0 |
| [D] | 97.5 ± 7.0 | 89.5 ± 11.5 | 108.2 ± 12.1 | 90.6 ± 23.8 |

### Frequency-dependent modulation at physiological temperature (36°C)

At 36°C, the overall trajectory of CAP changes during stimulus trains was similar to that observed at 22°C when responses were normalized to the first pulse in the train, however responses differ in magnitude (**Fig. 2**). Comparison of differences in frequency dependent amplitude changes between 5 Hz and 20 Hz at the last pulse displayed (pulse 9) were not significant at 22° but amplitude depression at 36° was significantly greater at 20Hz vs. 5Hz in both IGN and splanchnic populations (p<0.003 and p<0.021, respectively; paired t-test).

Analysis of fast- and slow-conducting components also revealed distinct temperature-dependent differences (**Fig. 4**). At 5 Hz (**Fig. 4A**), faster-conducting responses in both IGN and splanchnic nerves did not differ from baseline during the 20s train. In comparison, slower-conducting responses also showed no significant depression but exhibited post-train increases at 2 seconds (epoch D).

**Figure 4.**
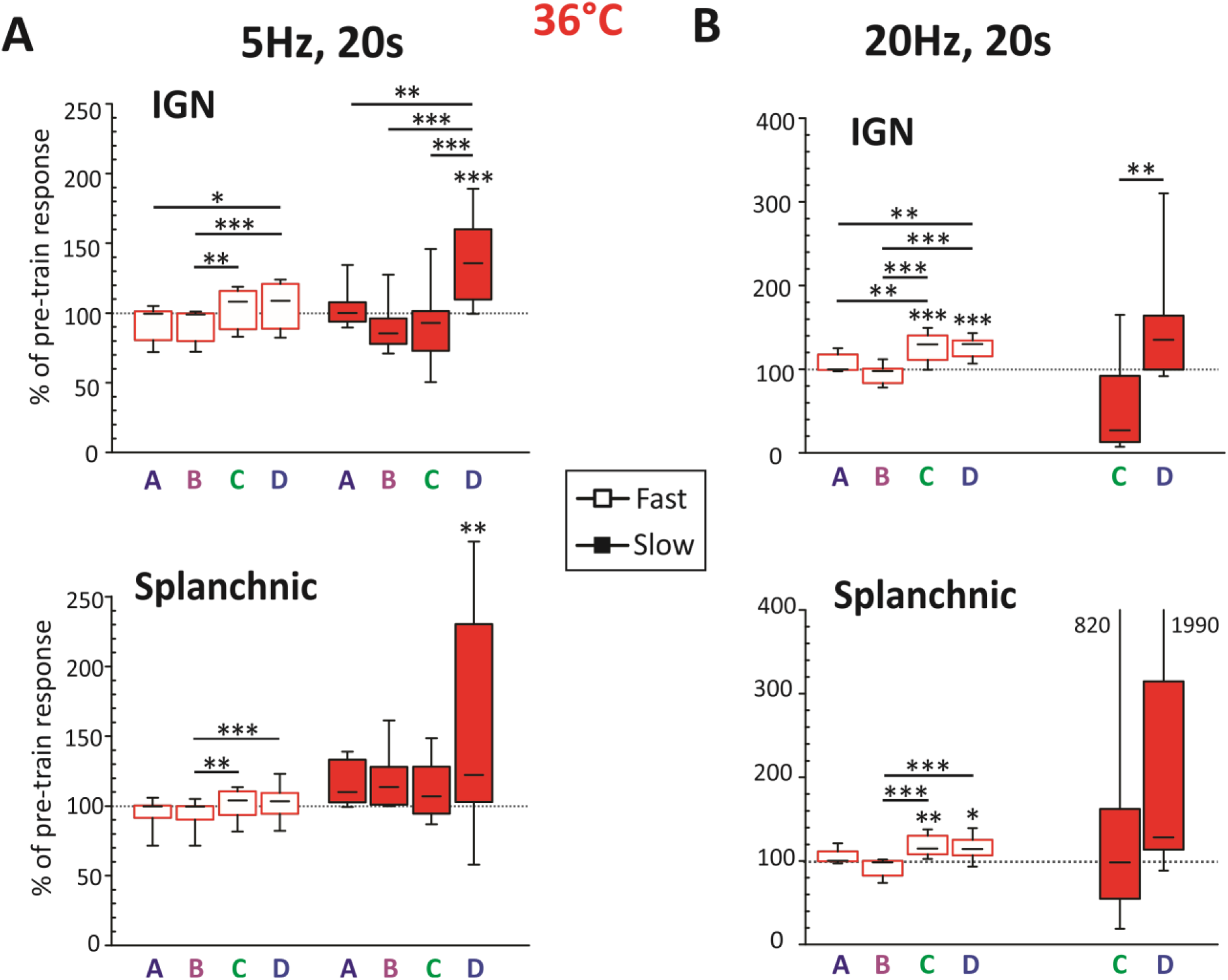
Effect of 5Hz and 20Hz trains on evoked response at 36°C. The rectified integral values for fast and slow CAP components are expressed as percentages of the pre-train pulse. **[A_1_]** 5 Hz. Fast axon CAPs in both IGN and splanchnic nerves were not different from baseline but events at the end of the train (C) and after conditioning (D) were larger that early responses in the train in both IGN and splanchnic. In comparison, the conditioned response (D) in slower conducting axons was significantly facilitated from baseline for both IGN and splanchnic nerves (138±11% and 152±28%, respectively). **[A_2_]** 20 Hz. Fast axons in both nerves exhibited a significant increase from baseline at the end of the train (D) which was sustained when examined 2s after the train (D). For fast IGN axons facilitation was 128±6% (C) and 127±4% (D) and for splanchnic, facilitation was 119±4% (C) and 116±5% (D) of baseline amplitude. In comparison, responses were more variable in slower conducting axons. The conditioned response (D) was facilitated in axons from the IGN nerve (150±25%). Statistical significance was assessed via repeated measures ANOVA and post hoc tests where relevant ([n=8]; *, p<0.05; **, p<0.01, **; and p<0.001, respectively).

At 20 Hz (**Fig. 4B**), faster-conducting responses increased during trains and remained elevated at 2s following the stimulus train. In the IGN, faster-conducting responses reached 127.7 ± 6.1% of baseline at the end of the train (p < 0.001) and remained elevated at 127.0 ± 4.3% at 2s post-train. Similar increases were observed in the splanchnic nerve. Slower-conducting responses were more variable and did not show consistent changes. A summary of results obtained at 36°C is provided in **Table 4**.

**Table 4.** 36C results of stimuli delivered at 5 & 20 Hz for 20 seconds (% of control). (*, p<0.05; **, p<0.01, **; and p<0.001, respectively)

| N=8 | Fast SPNs (0.58-3.4 m/s) |  | Slow SPNs (0.11-0.44 m/s) |  |
| --- | --- | --- | --- | --- |
|  | 5Hz | 20Hz | 5Hz | 20Hz |
| **IGN** |  |  |  |  |
| [A] | 93.9 ± 4.6 | 105.6 ± 4.1 | 103.2 ± 4.9 |  |
| [B] | 92.9 ± 4.3 | 95.3 ± 4.0 | 89.4 ± 6.2 |  |
| [C] | 104.5 ± 5.0 | 127.7 ± 6.1*** | 92.2 ± 9.9 | 53.9 ± 19.7 |
| [D] | 106.0 ± 5.6 | 127.0 ± 4.3*** | 138.4 ± 10.5*** | 149.7 ± 24.8 |
| **Spl.** |  |  |  |  |
| [A] | 95.7 ± 3.8 | 104.3 ± 3.1 | 116.2 ± 5.5 |  |
| [B] | 92.5 ± 3.8 | 93.3 ± 3.8 | 118.0 ± 7.5 |  |
| [C] | 101.8 ± 3.8 | 118.6 ± 4.4** | 111.0 ± 7.3 | 182.1 ± 93.4 |
| [D] | 103.0 ± 4.3 | 116.1 ± 4.9* | 151.6 ± 28.3* | 385.6 ± 231.3 |

### Interaction between temperature and activation history

Comparisons across temperatures indicate that frequency-dependent effects on CAP magnitude differ between axonal populations. At 22°C, slower-conducting components exhibited pronounced depression during repetitive stimulation, whereas at 36°C these components were reduced in magnitude under baseline conditions compared to 22°C (**Fig. 1E**) and failed to demonstrate statistically significant depression during trains. In contrast, faster-conducting components were largely preserved at 36°C (**Fig. 1E**) and exhibited stable or increased responses during high-frequency stimulation. Regardless of whether post train responses measured two seconds later were significantly changed, all responses consistently returned to baseline within the 60-second inter-episode window, indicative of shorter-term plasticity.

## DISCUSSION

Classical views assume SPN axons reliably relay centrally generated activity to postganglionic targets. In a separate study, we reported that there is a high recruitment variability at 22°C indicating this assumption does not hold (Halder et al., 2026). The present study shows that recruitment is further compromised at normal body T°. At 36°C, the population compound action potential (CAP) was reduced by approximately 50% relative to room temperature, with preferential loss of slower-conducting axonal components. These observations indicate a substantial reduction in effective axonal recruitment at physiological temperature and suggest that spike propagation along SPN axons is highly vulnerable to temperature-dependent disruption. Given that prior studies examining multisegmental sympathetic transmission were conducted at room temperature (Blackman and Purves, 1969; Nja and Purves, 1977; Lichtman et al., 1979, 1980; Purves and Lichtman, 1980), these findings raise the possibility that the extent of SPN influence across paravertebral ganglia has been overestimated.

The magnitude of the reduction in CAP amplitude at physiological temperature is most consistent with a loss of propagating spikes across a substantial fraction of SPN axons. After temperature-dependent changes in conduction velocity and spike waveform were accounted for through scaling procedures, the observed decrease in response magnitude greatly exceeded that expected from such biophysical effects alone. This suggests that a significant subset of axons fails to sustain action potential propagation at higher temperatures. This interpretation must be considered alongside possible or alternative explanations. Increased temperature may elevate recruitment thresholds, alter membrane excitability, or engage temperature-sensitive conductances that reduce the safety factor for propagation. Observations of lack of change in maximal recruitment at the stimulation intensity in which studies were undertaken ruled out reduced recruitment. The observed membrane hyperpolarization with warming is consistent with increased potassium conductance, which would both reduce excitability and attenuate electrotonic spread. Thus, the data strongly supports temperature-dependent reductions in conduction reliability versus altered recruitment threshold.

### Effects of temperature

To determine the impact of T° on spike propagation in SPNs, we compared volleys at room and normal body T°. At 36°C, both fast- and slow-conducting SPN axons experienced a T°-sensitive conduction block, failing at rates between 24-50%. Given the high variability in axonal recruitment observed in slow-conducting SPNs at 22°C (Halder et al., 2026), it is not surprising that these slow-conducting SPNs failed more frequently at 36°C compared to their faster-conducting counterparts. While the higher T°C promoted conduction block in all SPN axon populations, slower-conducting IGN axons were particularly susceptible. As the IGN contains branching axons, a putative mechanism is branch point failure.

### Temperature-sensitive leak channels and spike fidelity

RNA sequencing shows that the two-pore-domain potassium (K_2_P) channel TREK-1 (KCNK2), is present in thoracic SPNs (Alkaslasi et al., 2021). TREK-1 channel activity increases with increasing T°, with peak thermal responsiveness from 32-37°C, with ∼7-fold increase in current amplitude for every 10°C, indicating that physiological temperature variations can substantially influence its activity (Maingret et al., 2000). Increased TREK-1 channel activity may compromise spike conduction through two distinct mechanisms. (i) Hyperpolarization towards E_K_ would necessitate a larger depolarization for spike recruitment. (ii) An increased membrane conductance would reduce passive voltage amplitude and propagation space constant for regenerative spiking by downstream Na_V_ channels. Both hyperpolarization and increased membrane conductance would be expected to reduce the safety factor for spike propagation. In small-diameter axons, where electrotonic spread is already limited, such changes could readily precipitate conduction failure. While the present data are consistent with this mechanism, direct experimental manipulation of these channels will be required to establish their specific contribution.

### Small axon diameter preferential vulnerability to conduction failure

The preferential vulnerability of slower-conducting axons provides important insight into the structural determinants of this phenomenon. SPN axons are predominantly unmyelinated and extremely small in diameter, with many projecting across multiple paravertebral ganglia via branching pathways(Lewis and Burton, 1977; Halder et al., 2026). Smaller diameter axons are expected to operate with a reduced safety factor for propagation, rendering them particularly sensitive to perturbations in membrane conductance and spike waveform. Consistent with this, the greatest reductions in CAP magnitude were observed in slower-conducting populations, which likely correspond to the smallest caliber axons. Moreover, differences between interganglionic and splanchnic recordings suggest that axonal geometry contributes to conduction vulnerability. Interganglionic pathways, which include extensive branching within the sympathetic chain, exhibited greater temperature-dependent loss than splanchnic projections. These findings are consistent with the well-established susceptibility of axonal branch points to conduction failure, particularly under conditions that reduce available depolarizing current or increase membrane leak (Levy, 1980; Debanne et al., 2011). Together, these results identify axonal diameter and branching architecture as key determinants of temperature-sensitive conduction reliability in SPNs.

### Relevance of a population of silent SPNs

The interaction between temperature and activation history further reveals that changes in population response reflect selective filtering of axonal subpopulations rather than uniform modulation of all fibers. At room temperature, slower-conducting axons were present and exhibited pronounced frequency-dependent depression during stimulus trains, consistent with activity-dependent hyperpolarization and reduced excitability described in unmyelinated fibers (Renganathan et al., 2002; De Col et al., 2008; Obreja et al., 2010; Rama et al., 2018). In contrast, at physiological temperature, this vulnerable population was largely absent, and the remaining responses were dominated by faster-conducting axons that could sustain or even facilitate responses during higher-frequency stimulation. Thus, the apparent shift from depression at 22°C to facilitation at 36°C may not reflect a reversal of intrinsic frequency-following properties, but rather a change in the composition of the active axonal population. Temperature therefore acts as a filter that selectively removes axons with low conduction safety factor, leaving a more resilient subset that can support higher-frequency transmission. Relating the present data at 36°C to that obtained *in vivo*, would suggest that SPN signaling capacity would be highly compromised. Indeed, our results could contribute to why 40–70% of SPNs appear to be silent (lack ongoing or reflex activity) (Jänig, 2022).

### Changes in core body temperature may influence SPN recruitment

That SPN conduction undergoes more failures at 36°C than at 22°C may be physiologically important to differences in autonomic signaling. For example, while core body temperature range for humans and mice are similar (Reitman, 2018), mice are heterothermic that can enter hypometabolic torpor where core body temperature can reach 20°C (Geiser, 2004; Hrvatin et al., 2020). Given that mice exhibit large physiological fluctuations in body temperature during torpor, temperature-dependent modulation of SPN conduction could potentially contribute to adaptive autonomic regulation. For example, as reduced temperatures promote conduction, this may adaptively strengthen sympathetic drive to cutaneous vasoconstrictors for heat retention and to brown adipose tissue for heat generation through non-shivering thermogenesis (Enerbäck, 2009; Madden and Morrison, 2016; Francois et al., 2019). Additional studies would benefit from more detailed analysis of the effects of smaller temperature changes on the spike conduction in functional subpopulations of thoracic SPNs.

### Implications for sympathetic output regulation

Rather than functioning as a purely reliable transmission system, SPN axons appear to operate with reduced conduction safety margins, such that small changes in temperature or activation history can significantly alter the number and identity of axons that successfully propagate spikes. This introduces a mechanism of presynaptic gain control based on conduction reliability, whereby the effective size of the recruited postganglionic population can be dynamically regulated. As depression is frequency dependent in these smaller diameter axons, they are best suited to convey low-frequency population encoded responses to their postganglionic target neurons. Given this, one target population may be the smaller NPY-expressing vasomotor postganglionic neurons (Furlan et al., 2016) as observations in human microneurography show that individual postganglionic sympathetic neurons are recruited probabilistically with high variability in individual axonal recruitment with each baroreflex across cardiac cycles (Macefield and Wallin, 1999, 2018), suggesting that variability in sympathetic output may arise, in part, from constraints at the level of axonal conduction.

## Limitations

First, conclusions regarding conduction failure are inferred from population CAP measurements rather than direct recordings from individual axons. Although scaling procedures were used to account for temperature-dependent changes in spike waveform and conduction velocity, it remains possible that loss of axonal excitability rather than conduction failure contributed to the observed reductions in response magnitude. Second, classification of axonal populations was based on conduction latency rather than direct morphological identification, and thus the relationship between conduction velocity, axon diameter, and branching pattern remains inferential. Third, experiments were performed in an *ex vivo* preparation, and while this allowed precise control of temperature and stimulation parameters, it may not fully capture the complexity of in vivo modulatory influences. Finally, mice are nocturnal and experiments were undertaken during daytime. As ion channel expression/activity undergoes diurnal variation to reduce excitability during inactive periods (Itri et al., 2010; Colwell, 2011; Steponenaite et al., 2024), obtained results may reflect a time where recruitment is minimized.

### Summary

The present findings indicate that SPN axons operate with limited conduction reserve that is strongly modulated by temperature and activation history. Preferential vulnerability of slower-conducting, likely small-diameter and branching axons indicates that conduction failure represents a significant determinant of effective sympathetic output as a decrease in the magnitude of population responses would lead to diminished synaptic activity on postganglionic neurons. These results identify axonal conduction reliability as a physiologically regulated variable that can shape the strength and distribution of autonomic signaling.

## Acknowledgements

This research was funded by NIH NS121850, NS102871

